# PNPLA3^I148M^ is a novel regulator of bone mass independent of MASLD

**DOI:** 10.64898/2026.03.13.711662

**Authors:** Galen M Goldscheitter, Mulugeta Seneshaw, Faridoddin Mirshahi, Morgan B Summerlin, Allison C Ip, Austin H Coelho, Ronnie Li, Damian C Genetos, Arun J Sanyal, Henry J Donahue

## Abstract

Metabolic-dysfunction associated steatotic liver disease (MASLD) is the most common chronic liver disease. Fracture risk is increased among people with MASLD, however, the genetic contribution to risk is undetermined. PNPLA3^I148M^ is a common SNP which accounts for most MASLD heritability and increases MASLD morbidity and mortality. However, PNPLA3^I148M^ impact on bone is unexplored. To bridge this gap, we used a validated murine model of MASLD (DIAMOND mice) which received human PNPLA3 transgenes via adeno-associated vector serotype 8 (AAV8) and assessed bone morphology, cellularity, and transcriptomics. PNPLA3^I148M^ was expressed in bone and associated with bone loss, decreased bone formation, increased bone resorption, and increased bone marrow adiposity. PNPLA3^I148M^ reprogrammed the transcriptome in bone, enriching expression of pathways associated with fatty acid metabolism and hampering bone turnover. Notably, these findings occurred in the absence of MASLD. These findings suggest PNPLA3^I148M^ possesses an intrinsic deleterious skeletal role.

## INTRODUCTION

Metabolic dysfunction-associated steatotic liver disease is ∼38% prevalent globally [1]. It is a heterogenous disease producing increased liver, cardiovascular, and endocrine morbidity and mortality [2]. We recently identified increased fracture risk among a large cohort of people with MASLD, adding an adverse outcome in this population [3]. Fractures, especially among older adults, increase morbidity and mortality [4]. The genetic basis of increased fracture risk in MASLD is unknown and presents a major obstacle to development of safe, effective treatments.

Patatin-like phospholipase domain containing 3 (PNPLA3) is the first identified genetic determinant of MASLD risk and severity [5]. PNPLA3 *rs738409* C>G missense mutation encodes an isoleucine to methionine conversion at codon 148 (PNPLA3^I148M^), a gain-of-function variant leading to increased ballooning degradation, inflammation, and fibrosis [6]. PNPLA3^I148M^ is 17-49% prevalent with highest frequency in Hispanic populations, the subpopulation with greatest MASLD prevalence [5]. PNPLA3^I148M^ is the principal driver of MASLD heritability, however, its contribution to MASLD-associated skeletal fragility is unknown [7]. It is critical to address this gap to mitigate MASLD-associated fracture morbidity.

Bone is a complex multicellular tissue providing structure and protection in vertebrate organisms. It supports metabolic, hematopoietic, immune, and endocrine functions via metabolism of its mineralized matrix and via the resident bone marrow [8]. Skeletal fragility arises when bone metabolism is dysregulated by genetic or environmental insults to myriad functions of bone [9]. In MASLD, metabolic syndrome, dysregulated hepatokine secretion, and impaired vitamin D metabolism may all contribute to bone loss [3], [10]. A single report showed PNPLA3^I148M^ associates with low bone mineral density in adolescents with histologically confirmed MASLD, though MASLD severity confounded this result [11]. Other work shows MASLD severity correlates with lower bone mineral density in another adolescent population [12]. The impact of PNPLA3^I148M^ on bone is therefore unclear, a major gap addressed in this work.

While PNPLA3^I148M^ is a common variant well classified in hepatic steatosis, its role in bone metabolism is unstudied. Sequencing datasets show PNPLA3 expression in bone marrow, however its presence in mineralized bone compartment is undetermined [13]. Given the increased risk of fracture among people with MASLD population, and high prevalence of PNPLA3^I148M^, it is imperative to characterize the role of the variant on bone metabolism to abrogate increased fracture morbidity and mortality.

We recently established DIAMOND mice, a validated preclinical model of MASLD, as a model of MASLD-associated skeletal fragility [3], [10]. Additionally, we previously leveraged this model to investigate gene-environment interactions accelerating MASLD progression with PNPLA3^I148M^ [14]. In the current study, we combine this basis to explore PNPLA3^I148M^ impact on bone. The first aim of this study was to evaluate the impact of PNPLA3^I148M^ overexpression on bone morphology and strength. Secondly, we sought to investigate dysregulation of bone metabolism by PNPLA3^I148M^ at the cellular level. The third aim was to explore transcriptomic reprogramming in bone by PNPLA3^I148M^.

## METHODS

### Animal Studies

Sixty 8-week-old male DIAMOND mice (an isogenic cross between C57BL6/J and 129S1/SvImJ mice) were randomized to 3 adeno-associated virus serotype-8 (AAV8) vectors containing: luciferase and a scrambled sequence (Luc), luciferase and human PNPLA3 wild-type (PNPLA3^WT^), or luciferase and human PNPLA3^I148M^ (PNPLA3^I148M^). Vectors were obtained from the University of Maryland virology core. Mice were further randomized to chow diet and normal water (CD/NW, Teklad.7012) or high-fat diet and sugar water (HFD/SW, Teklad.88137, 42% calories from fat). Mice were maintained on their diets in standard house conditions (12h light-dark cycle, 22-24 °C, 55% relative humidity) for 36 weeks after vector delivery and diet initiation. A supplementary vector bolus was delivered 16 weeks following first bolus (**Figure 1**). Animals were humanely euthanized via CO_2_ asphyxiation.

**Figure 1:**
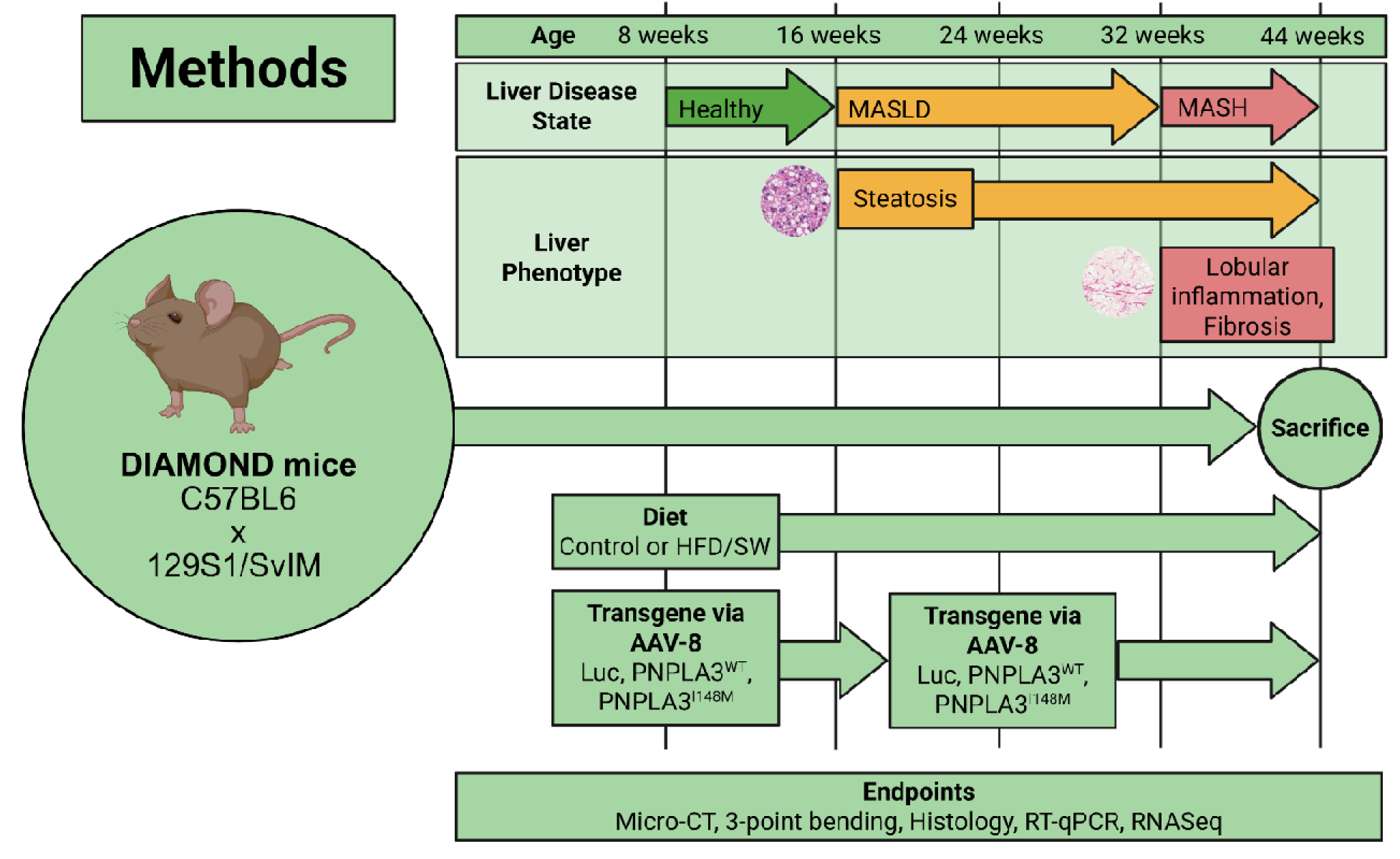
Design of animal experiments and corresponding hepatic phenotype at each timepoint for animals on HFD/SW.

### Generation of viral vectors

Vectors were generated as previously described [14]. AAV8s were prepared by the Viral Vector Core at the University of Massachusetts Medical School. Human wild-type and I148M variant PNPLA3 were synthesized and cloned into a pUC57 shuttle plasmid (Genewiz LLC), with 5’ MluI sites and 3’ SalI sites. The sequences were subcloned into MluI and SalI sites of a liver-specific thyroxine binding globulin-containing AAV8 plasmid, with luciferase as a control.

### Tibia geometric properties

Tibias were excised, wrapped in PBS-soaked gauze, and stored at−80 °C. Bones were embedded in a 3D-printed mold in a 1% (m/v) agarose matrix. Bones were imaged using a benchtop micro-computed tomography (µCT) scanner (Bruker SkyScan 1276) with the following parameters: 0.5 mm aluminum filter, 200 kV detector potential, 60 µA detector current, 730 ms integration time, 0.5° rotation step. µCT scans were reconstructed, aligned, segmented, and analyzed using the Bruker software suite (Bruker NRecon, DataViewer, CTAn). Mid-diaphyseal cortical bone was analyzed in 180 µm-long regions of interest (ROIs). Trabecular bone was analyzed at distal metaphysis and distal epiphysis: 540 µm-long metaphyseal ROIs were established beginning 200 µm proximal to the distal epiphyseal plate, while 400 µm-long epiphyseal ROIs were established beginning at the distal epiphyseal plate. Bone morphometry was analyzed as previously described [10]. In both metaphyseal and epiphyseal regions, trabecular bone was manually segmented by a blinded evaluator aided by software-driven interpolation.

### Tibia mechanical properties

Tibias were excised, thoroughly cleaned of all soft tissue, wrapped in PBS-soaked gauze, and stored at−80 °C. Tibias were µCT scanned before mechanical testing for assessment of geometry-independent parameters. Mechanical properties were assessed via 3-point bending (100 lbf load cell, Bose ElectroForce 3200). Tibias were brought to room temperature and placed on a 10 mm support span with the anteromedial surface in tension and the apex of primary curvature centered between the supports. A mover was centered between the supports, aligned with the apex of primary curvature, and driven downwards at 1 mm/min until failure occurred by transverse fracture. Load and displacement data were captured at 10 Hz. Quantification and analysis of mechanical testing parameters was performed using previously described methods [15].

### Histologic sample preparation

Femurs were dissected—leaving 2-3 mm of muscle in place—and fixed in 10% neutral buffered formalin for 48 hours at 4 °C with gentle agitation. Formalin-fixed tissues were embedded in paraffin using standard methods. 5 µm sections were prepared using a benchtop microtome and adhered to positively charged slides in preparation for histologic staining. After all staining procedures, whole slide areas were scanned using a 40X objective (Akoya Biosciences PhenoImager HT). Analysis of all histologic samples was performed by blinded evaluators using QuPath (v0.6.0) [16].

### Osteocyte density

Femur sections were prepared as described above, deparaffinized, and stained with Mayer’s hematoxylin and eosin using standard methods. 1 mm ROIs were established at mid-diaphyses. Osteocytes were defined as eosin-positive cells occupying intact lacunae within mid-diaphyseal cortices. An unsupervised blinded classifier was trained using historical data and was used to count osteocytes and empty lacunae (QuPath v0.6.0) [16]. Osteocyte number and empty lacunae number were normalized to cortical bone area.

### Osteoblast number, mineralizing surface, and activity

Femur sections were deparaffinized and prepared using immunohistochemical methods. Samples were blocked with normal goat serum, peroxidases were blocked using 3% H_2_O_2_, IgG against murine Sp7/Osterix was applied, and detection was performed via a biotinylated secondary antibody conjugated to a streptavidin-HRP complex and 3,3’-diaminobenzidine. Slides were scanned as described above. A 1 mm ROI was established in mid-diaphyses. Osteoblasts were identified as Sp7/Osx^+^ cuboidal cells attached to bone surfaces. Osteoblast number, and their mineralizing surface, were normalized to the total bone surface within the ROI

### Osteoclast number, mineralizing surface, and activity

Femur sections were deparaffinized and stained using a tartrate-resistant acid phosphatase (TRAP)-based enzymatic stain. 1.5 mm ROIs established in distal metaphyses. Osteoclasts were identified as TRAP^+^ multinuclear cells attached to bone surfaces. Osteoclast number was counted and normalized to the total bone surface within the ROI. Osteoclast surface was measured as the surface of the resorption pit associated with each osteoclast and normalized to total bone surface.

### Adipocyte number and area

Adipocytes were assessed via immunohistochemical staining. Deparaffinized sections were stained, as described above, using a primary antibody against perilipin-1 (PLIN-1). A 1.5 mm ROI was established in distal metaphyses. Adipocytes were considered PLIN-1^+^ cells with central clearing. The total number, and area covered by adipocytes, were summed and normalized to the total tissue area within the ROI.

### mRNA isolation and reverse-transcription quantitative polymerase chain reaction

Femurs and livers were isolated for gene expression analysis. Proximal and distal heads of the femurs were removed, bone marrow was removed via centrifugation, and periosteum was scraped away. Bone and liver tissues were fixed overnight in RNAlater (Invitrogen AM7020) at 4°C. The following morning, RNAlater was aspirated off, and samples were stored at−80 °C. Bones were crushed in liquid nitrogen using a mortar and pestle. Crushed bone and liver tissues were lysed in RLT-plus lysis buffer (QIAGEN 1053393) and homogenized using a bead mill (ThermoFisher 15-340-164). RNA was isolated using RNeasy Plus Micro Kits (QIAGEN 74034). Total RNA concentration and A260/280 ratio were measured on a NanoDrop Lite spectrophotometer (ThermoFisher NDNDLUSCAN). Complementary DNA was synthesized using iScript reverse transcriptase and random primers (BioRad 1708891). Gene expression was measured by real-time PCR via SYBR green fluorescence on a C1000 Touch thermocycler (BioRad 1851148) with CFX96 optical reaction module (BioRad 1845097). Human *PNPLA3* expression was measured in bone using the primer pair qHsaCID0007552 (BioRad). Gene expression was normalized to murine *Actb* using the primer pair qMmuCED0027505 (BioRad).

### mRNA sequencing and alignment

Messenger RNA quality was assessed using an Agilent 2100 Bioanalyzer. Samples with RNA integrity number ≥7.0 were sequenced by NovoGene (Sacramento, CA), in bulk, using an Illumina NovaSeqX Plus. Raw sequence data was adapter trimmed, aligned, and annotated using Nextflow software (nf-core/rnaseq v3.14.0). Sequence data was mapped to mouse genome mm10 using STAR (v2.7.11b) and annotated to GENCODE vM25. Quality control was performed using RseQC, Qualimap, and principal component analysis within nf-core/rnaseq. Correlation plots were also created by calculating Pearson’s correlation coefficient and used to generate heatmaps demonstrating alignment of gene expression within experimental groups.

### Differential expression analysis

Differential expression analysis was performed using DESeq2 (v1.44.0). Differential expression was compared between conditions using Wald testing, with multiple comparisons controlled using the Benjamini-Hochberg method. False discovery rate threshold (p_adj_) for significance was set at 0.05 with an absolute log_2_ fold change (LFC) greater than 1. Genes without ≥10 reads in ≥50% of samples were considered not expressed and were excluded from the analysis.

### Biological process and molecular pathway enrichment

We performed gene ontology (GO) analysis and gene-set enrichment analysis (GSEA) to assess the impact of differential gene expression on molecular pathways and biological functions. GO terms and associated genes were obtained from the Molecular Signatures Database (MSigDB, *mus musculus,* v2025.1). We performed an initial analysis of the orthology-mapped hallmark gene sets (MH, 50 gene sets) using fast gene set enrichment analysis (fgsea v1.30.0). To increase resolution, this was followed by GSEA of all biological process GO terms (GO:BP, 7583 gene sets). Guided by results of the hallmark GSEA, we further restricted the analysis to GOBP terms associated with adipogenesis and osteogenesis (See Supplemental). GO terms and pathway activities are visualized as normalized enrichment score with color intensity denoting associated family-wise error rate-adjusted p value.

### Statistical analysis

For non-sequencing outcomes, statistical comparisons and graphing were performed in GraphPad Prism (v10.5.0), with outlier detection and removal in MATLAB (v2024a). All data are presented as mean ± standard error of the mean. Normality of residuals and equality of variance were tested via Anderson-Darling and Bartlett methods, respectively. Intergroup differences among normally distributed and homoscedastic outcomes were assessed using ordinary one-way ANOVA, when heteroscedastic, Welch’s ANOVA was used. In these cases, *post-hoc* testing, with family-wise error rate control, was performed using the Tukey procedure. When samples were not normally distributed, the Kruskal-Wallis test was used to assess differences. A threshold two-tailed p-value of 0.05 was used for all comparisons. Animal studies were powered to detect decreased cortical thickness after 36 weeks of HFD/SW exposure based on previous work with α = 0.05 and β = 0.20 [3], [10]. The statistical analysis plan for RNA sequencing is detailed above. All bioinformatics analysis was performed using R (v.4.4.1).

## RESULTS

### Physiologic and skeletal changes in DIAMOND mice fed HFD/SW vs CD/NW

After 36 weeks of HFD/SW feeding, DIAMOND mice became obese and developed hepatomegaly compared to those fed CD/NW (**Supp. Fig. 1A)**. PNPLA3^WT^ mice fed HFD/SW lost cortical thickness (Ct.Th) compared to PNPLA3^WT^ mice fed CD/NW (**Supp. Fig. 1B**). PNPLA3^I148M^ mice gain epiphyseal trabecular bone volume fraction (BV/TV), trabecular number (Tb.N) and decrease trabecular spacing (Tb.Sp) on HFD/SW vs CD/NW (**Supp. Fig. 1C**). PNPLA3^WT^ and PNPLA3^I148M^, but not Luc mice, gain metaphyseal trabecular bone volume (BV/TV) and trabecular number (Tb.N) (**Supp. Fig. 1D**). Luc and PNPLA3^WT^ mice lose trabecular thickness on HFD/SW vs CD/NW (**Supp. Fig. 1D**). All genotypes decrease trabecular spacing on HFD/SW vs CD/NW (**Supp. Fig. 1D**). In 3-point bending of tibias, Luc and PNPLA3^WT^ mice fed HFD/SW vs CD/NW lose ultimate load borne before failure and load at yield (**Supp. Fig. 2**). Luc mice fed HFD/SW vs CD/NW lose stiffness and work to fracture (**Supp. Fig. 2**) Young’s Modulus increased among PNPLA3^WT^ mice fed HFD/SW vs CD/NW (**Supp. Fig. 2**). Finally, ultimate deformation decreased among PNPLA3^I148M^ mice fed HFD/SW vs CD/NW (**Supp. Fig. 2**). Changes observed among PNPLA3^WT^ and Luc mice on HFD/SW vs. CD/NW aligned with those in our previously published work [3], [10]. Therefore, the remainder of this study focused on differences between genotypes independent of MASLD, and CD/NW data are presented from here forward.

**Figure 2:**
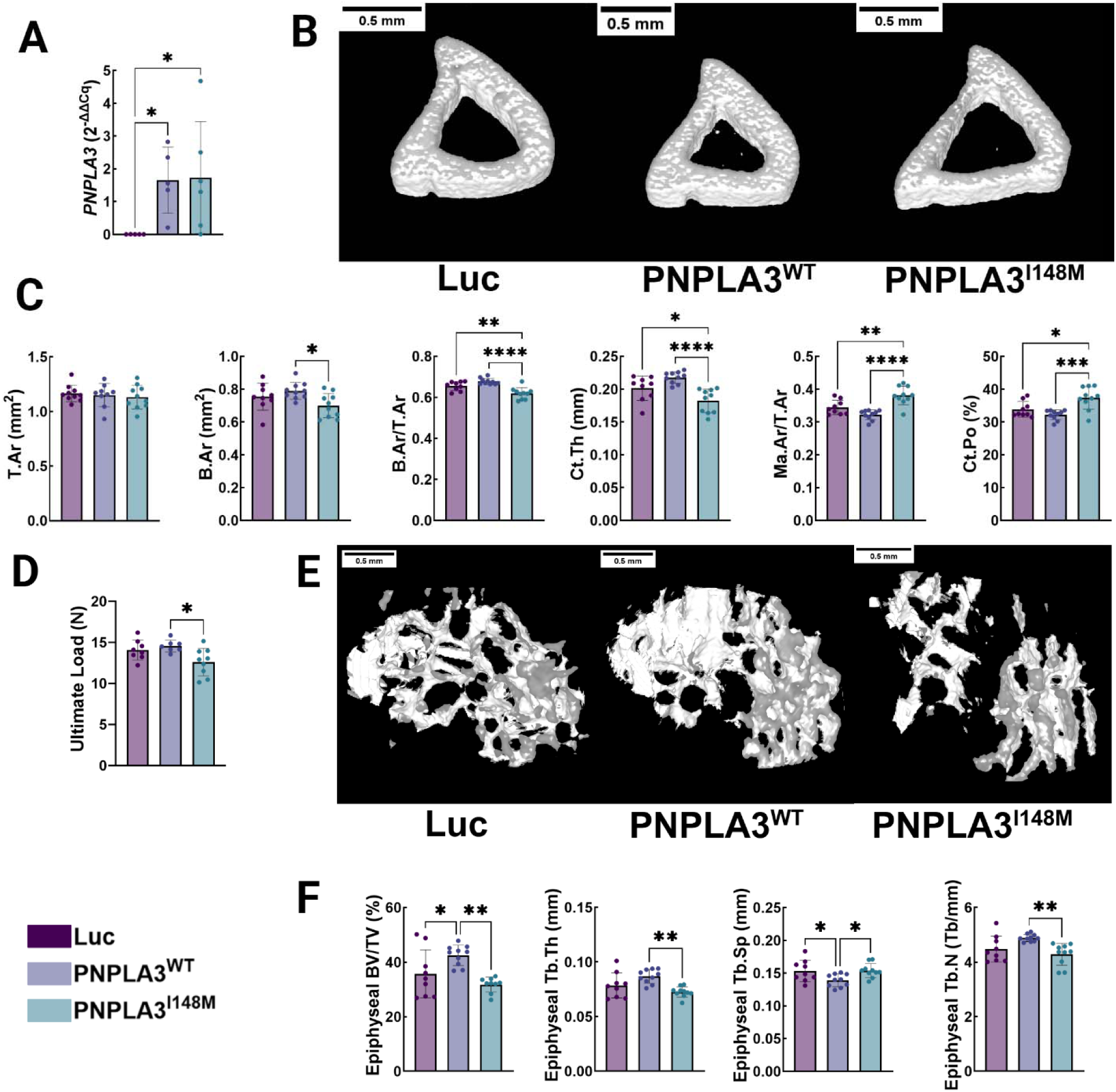
Human PNPLA3 is expressed in bone and deleterious bone outcomes are seen in PNPLA3^I148M^ mice on CD/NW. (**A**) Human PNPLA3 expression in bone via RT-qPCR. (**B**) 3-dimensional reconstructions of tibial cortical bone via µCT. (**C**) Tibial cortical bone morphology. (**D**) Ultimate load achieved in 3-point bending. (**E**) 3-dimensional reconstructions of tibial epiphyseal bone via µCT. (**F**) Tibial epiphyseal trabecular bone morphology. (* p<0.05, ** p<0.01, *** p<0.001, ****p<0.0001)

### Human PNPLA3 delivered via AAV8 was expressed in bone

Human *PNPLA3* was expressed among DIAMOND mice treated with AAV8-PNPLA3^WT^ and AAV8-PNPLA3^I148M^ constructs. Meanwhile, no human *PNPLA3* was expressed among mice treated with AAV8-Luc construct (**Fig. 2A**).

### Human PNPLA3^I148M^ associated with cortical and trabecular bone loss without MASLD

36 weeks following first exposure to AAV8 constructs, PNPLA3^I148M^ DIAMOND mice lost cortical bone in the mid-diaphysis and trabecular bone in the proximal epiphysis of tibias. PNPLA3^I148M^ mice lost cortical thickness (Ct.Th) compared to both Luc and PNPLA3^WT^ mice. **(Fig. 2B & 2C)** The average bone area (B.Ar) of PNPLA3^I148M^ mice decreased compared to PNPLA3^WT^ mice, but not Luc mice (**Fig. 2B & 2C**). When correcting for tissue area (T.Ar), the relative bone area (B.Ar/T.Ar) among PNPLA3^I148M^ mice decreased compared to both PNPLA3^WT^ and Luc mice; we observed concomitant increased relative medullary cavity size (Ma.Ar/T.Ar) in PNPLA3^I148M^ mice compared to both PNPLA3^WT^ and Luc mice (**Fig. 2B & 2C**). Decreased cortical thickness among PNPLA3^I148M^ was further reflected via increased relative medullary cavity size (Ma.Ar/T.Ar) compared to both PNPLA3^WT^ and Luc mice (**Fig. 2B & 2C**). Cortical porosity increased among PNPLA3^I148M^ mice vs PNPLA3^WT^ and Luc mice (**Fig. 2B & 2C)**. Ultimate load borne in 3-point bending was decreased among PNPLA3^I148M^ mice compared to PNPLA3^WT^ mice, with no changes observed in Luc mice (**Fig. 2D**). Other markers of mechanical integrity were unaffected (**Supp. Fig. 2**). Within the proximal epiphysis of the tibia, PNPLA3^WT^ mice exhibited higher relative bone volume (BV/TV) than PNPLA3^I148M^ and Luc mice (**Fig. 2E & 2F**). Trabecular thickness and number were lowered in PNPLA3^I148M^ mice compared to PNPLA3^WT^ mice, with no differences observed among Luc mice and other groups (**Fig. 2E & 2F**). Trabecular spacing increased among PNPLA3^I148M^ and Luc mice compared to PNPLA3^WT^ mice (**Fig. 2E & 2F**). No differences were observed in metaphyseal trabecular bone on CD/NW (**Supp. Fig. 1D**).

### Cellular bone metabolism indicators are dysregulated with PNPLA3^I148M^ without MASLD

Osteoblast formation and osteocyte survival were impaired, meanwhile osteoclast formation and bone marrow adipogenesis were increased among PNPLA3^I148M^ mice (**Fig. 3A**). Osteoblast number relative to medullary bone surface (Ob.N/BS) decreased among PNPLA3^I148M^ mice compared to PNPLA3^WT^ and Luc mice (**Fig. 3B**); correspondingly, osteocyte number normalized to cortical bone area (Ot.N/B.Ar) also decreased in PNPLA3^I148M^ mice compared to Luc mice (**Fig. 3C**). Surface covered by osteoclasts relative to total bone surface (Oc.S/BS) increased among PNPLA3^I148M^ mice compared to PNPLA3^WT^ and Luc mice (**Fig. 3D**). Adipocyte number and Adipocyte area in bone marrow normalized to medullary area (Adp.N/M.Ar) also increased among PNPLA3^I148M^ mice compared to Luc mice (**Fig. 3E**).

**Figure 3:**
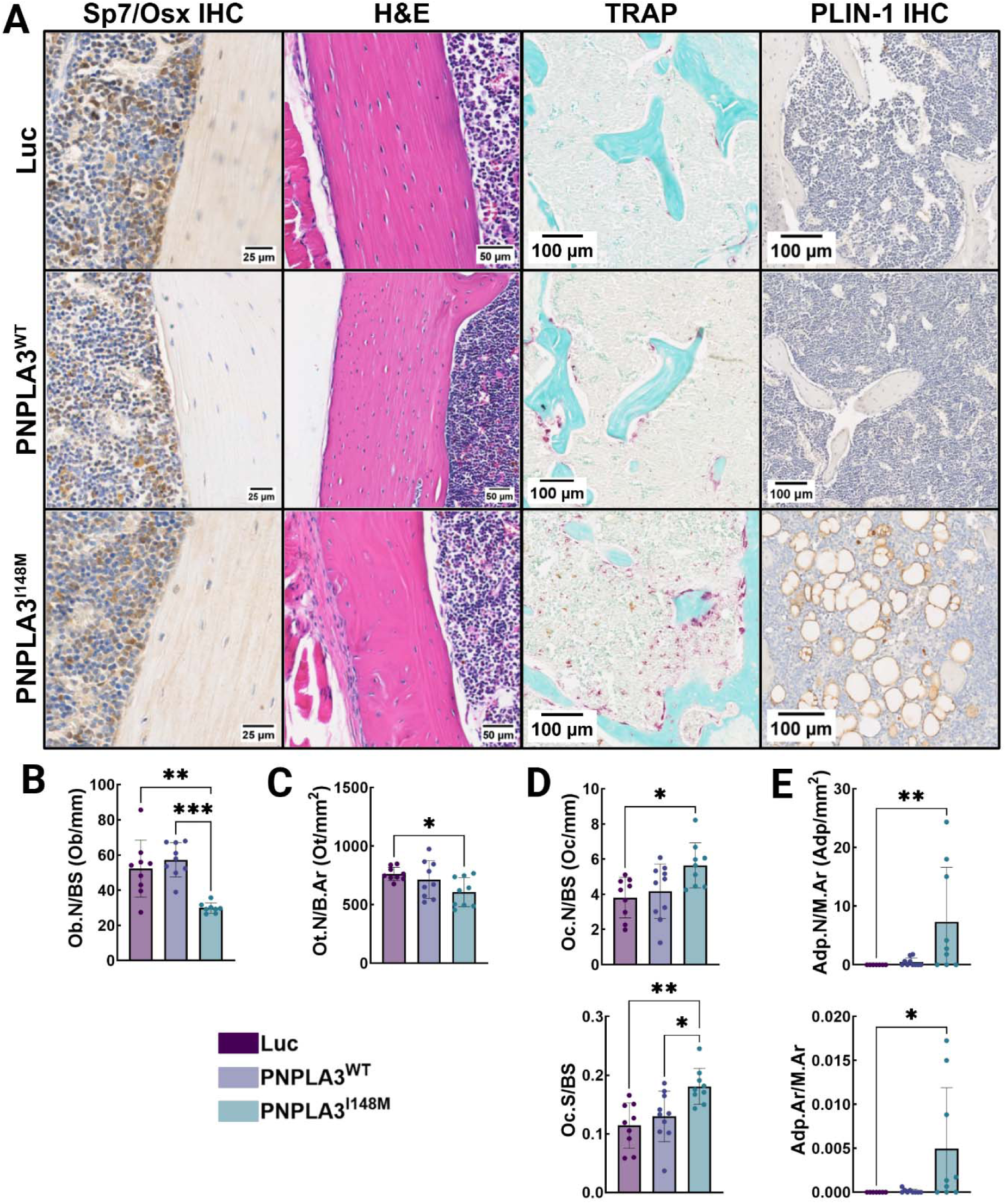
PNPLA3^I148M^ mice lose osteoblasts, osteocytes, and gain osteoclasts and bone marrow adipocytes on CD/NW. (**A**) Osteoblast identification via Sp7/Osx IHC stain, cortical H&E stain, osteoclast staining via TRAP, and adipocyte identification via PLIN-1 IHC stain. (**B**) Osteoblast number per bone surface. (**C**) osteocyte density per bone area. (**D**) Osteoclast number and density per bone surface. (**E**) Adipocyte number and area per marrow area. (* p<0.05, ** p<0.01, *** p<0.001)

### Bone metabolism and adipogenic pathways are reprogrammed with PNPLA3^I148M^

Unbiased bulk RNA sequencing was used to evaluate transcriptomic differences among genotypes within bone. Differential gene expression was compared between PNPLA3^WT^ and PNPLA3^I148M^ mice with quality control via PCA supported by counts heatmap (**Fig. 4A**). GSEA using hallmark gene sets revealed increased expression of Myc targets, fatty acid metabolism genes, and adipogenesis-associated genes, meanwhile inflammation (TNFα signaling via NF-κB, IL6 JAK-STAT signaling) were downregulated (**Fig. 4B**). Evaluating all biological process ontology via GSEA demonstrated increased electron transport, oxidative phosphorylation, and translation-associated gene expression (**Fig. 4C**). Adipo- and osteogenic subset analysis revealed increased lipid oxidation gene sets, meanwhile, bone resorption, remodeling, vitamin D metabolism, and triglyceride metabolism were inhibited (**Fig. 4D**). Increased *Lep* and *Plin5* expression were principal changes enriching increased beta oxidation ontology (**Fig. 4E, Supp. Fig. 5**). Meanwhile, decreased *Fcgr4* and *Il6* expression enriched inhibition of bone resorption ontology (**Fig. 4F, Supp. Fig. 6**) Changes between Luc and PNPLA3^I148M^ mice mirrored those seen between PNPLA3^WT^ and PNPLA3^I148M^ mice (**Supp. Fig. 3, 7, 8**). Differential gene expression analysis between Luc and PNPLA3^WT^ mice revealed few differential expressed genes, and failed to separate on PCA (**Supp. Fig. 4**).

**Figure 4:**
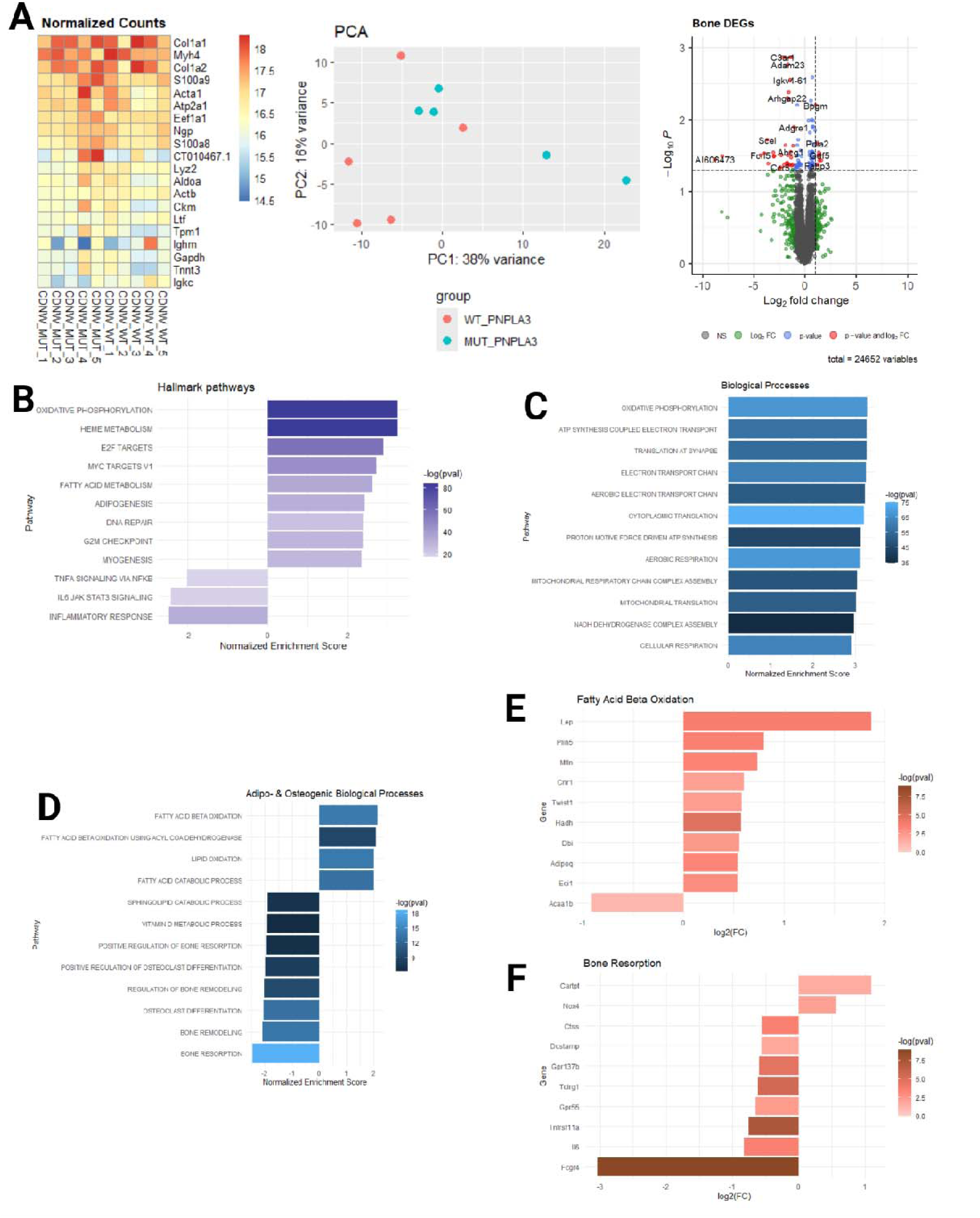
Metabolic, adipogenic, and osteogenic pathways are reprogrammed with PNPLA3^I148M^ vs PNPLA3^WT^. (**A**) Highest expressed genes, separation by PCA, and DEGs in bone from PNPLA3^I148M^ vs PNPLA3^WT^ mice. (**B**) Hallmark pathway, (**C**) biological process, and (**D**) adipo-and osteogenic biological process ontology pathway enrichment in PNPLA3^I148M^ mice compared to PNPLA3^WT^ mice. Individual gene contributions to changes in (**E**) beta oxidation and (**F**) bone resorption biological process ontology terms in bone from PNPLA3^I148M^ vs PNPLA3^WT^ mice

## DISCUSSION

MASLD associates with adverse skeletal outcomes. Previous work by our group and others described skeletal fragility in mice and humans with MASLD [3], [10], [17]. We therefore investigated the impact of PNPLA3^I148M^, a genetic driver of MASLD, on skeletal form and function. PNPLA3^I148M^ increases the odds of developing MASLD and increases morbidity and mortality among people with MASLD [5]. While studying interaction between hepatic PNPLA3^I148M^ and bone health in DIAMOND mice, we identified a direct deleterious effect of PNPLA3^I148M^ on bone. This exciting, serendipitous finding motivated diet-agnostic investigations into skeletal PNPLA3^I148M^ effects.

Human PNPLA3 transgenes were delivered via AAV8 vector system because the system exhibits high hepatocyte transduction selectivity [18]. To increase this selectivity of our model system, we employed a liver-specific thyroxine binding globulin promoter [14]. However, here we show expression of the transgene within the appendicular skeleton and subsequently leverage this finding to investigate the skeletal impact of PNPLA3^I148M^. To our knowledge, expression of AAV8-delivered transgenes in bone has not been reported by other authors.

Cortical bone comprises >90% of long bone mass and accounts for most of its fracture resistance [19]. Cortical volume and thickness decreased in PNPLA3^I148M^ mice, meanwhile, medullary cavity volume and cortical bone porosity increased. Overexpression of human PNPLA3^I148M^, but not PNPLA3^WT^ or Luc, therefore, associates with losses within the bone compartment which provides most failure resistance; functionally, bones from PNPLA3^I148M^ mice failed at lower ultimate load values in 3-point bending. In trabecular bone, PNPLA3^WT^ mice developed greater bone volume, thickness, number, and lower spacing than PNPLA3^I148M^ mice. However, trabecular bone volume was lower and trabecular spacing was higher among Luc mice than PNPLA3^WT^ mice, suggesting protection via PNPLA3^WT^ in trabecular bone. Indeed, no differences were observed in trabecular bone among Luc and PNPLA3^I148M^ mice. We show strong support for a deleterious impact of PNPLA3^I148M^ on cortical bone, suggesting increased fracture risk, and a possible protective role of PNPLA3^WT^ in trabecular bone.

Mineralized bone matrix is supported by three principal cell types: osteoblasts, osteocytes, and osteoclasts [20]. We investigated the role of PNPLA3^I148M^ on the distribution of these cells histologically. Both osteoblast and osteocyte density decreased in PNPLA3^I148M^ mice. Fewer osteoblasts mitigates new bone formation. Osteocytes are terminally differentiated osteoblasts, embedded in mineralized bone matrix. Decreased osteocyte density and dendritic network connectivity are typically seen in aging, associating with poor skeletal outcomes [21], [22]. Osteoclasts, meanwhile, were more abundant in PNPLA3^I148M^ mice, indicating an increase in bone catabolism. The combined effects of decreased formation and increased resorption synergistically accelerate bone loss. A potential explanation for these findings may be found in the bone marrow compartment, where marrow adiposity increased among PNPLA3^I148M^ mice. Increased bone marrow adiposity (BMA) is associated with decreased bone mineral density in humans [23], [24]. BMA and osteoblast presence are inversely related [25]. Further, increased adipokine secretion via increased adipocyte abundance may accelerate osteoclastogenesis and subsequently increased bone resorption [26], [27], [28]. While early osteoblasts rely heavily on glycolysis for energy production, both mature osteoblasts and adipocytes rely heavily on fatty acid β-oxidation [29]. Because PNPLA3 mobilizes fatty acids for β-oxidation, both osteoblasts an adipocytes may be more susceptible—relative to other cells in bone—to dysregulation via PNPLA3^I148M^ overexpression.

Analysis of the skeletal transcriptome comparing PNPLA3^WT^ and PNPLA3^I148M^ mice enabled identification of candidate mechanisms driving skeletal fragility among PNPLA3^I148M^ mice. Strongest signals in GSEA were oxidative phosphorylation and heme metabolism hallmark pathways. Oxidative phosphorylation and heme metabolism are mediators of osteoclastogenesis and may support increased osteoclast presence seen histologically [30]. However, in contrast to our histologic findings, osteoclast differentiation and bone resorption pathways decreased among PNPLA3^I148M^ mice compared to PNPLA3^WT^ mice. Increased BMA may account for this conflicting finding [25]. Adipocytes may replace other cell types, resulting in decreased relative osteoclast RNA representation among PNPLA3^I148M^ compared to PNPLA3^WT^ mice. This effect may also explain enrichment of pathways associated with beta oxidation and catabolism. Within a clonal population of adipocytes, increases in fatty acid catabolism would associate with decreased, not increased, adipocyte size [31]. However, the heterogenous cell population in our sequencing dataset cannot account for this. A single-cell sequencing study is needed to investigate this further. Hallmark pathways associated with Myc targets, fatty acid metabolism, and adipogenesis were upregulated, which we further investigated within biological process ontology terms. Within beta oxidation, we expect increased *Lep* (encodes leptin) to associate with skeletal fragility, likely by starving osteoblasts of energy needed for ossification [32], [33]. While osteoclasts were indeed increased, there was a concomitant decrease in osteoblasts. Osteoclast activity and osteoblast activity are necessarily coupled within the adult skeleton [34]. Impairment of osteoblast function by PNPLA3^I148M^ would therefore indirectly inhibit bone catabolism by osteoclasts. These findings suggest that PNPLA3^I148M^, by impairing lipid transport, may be directly antagonizing osteoblast function, driving a low-bone mass phenotype.

This study, like all others, has shortcomings. We only employed male mice because we previously demonstrated only male DIAMOND mice develop skeletal fragility with MASLD [10]. However, the sexual dimorphism of genetic determinants of skeletal fragility may differ from the environmental determinants and should be addressed in future work. This study was powered to detect differences in mechanical testing outcomes between DIAMOND on HFD/SW vs CD/NW at 48 weeks from previous studies. The cellular distribution of PNPLA3 overexpression could not be assessed with the methods employed. Finally, the PNPLA3 overexpression was not physiologic and there are meaningful differences between human and murine lipid metabolism [14], [35], [36].

In summary, the current study demonstrates an association between PNPLA3^I148M^ and skeletal fragility underpinned by decreased bone formation, increased bone resorption, and increased bone marrow adiposity. Remarkably, these findings occurred in the absence of high-fat diet feeding, and therefore in the absence of MASLD. These findings were further supported by reprogramming of bone metabolism and BMA-regulating networks. These findings provide insight into the skeletal biology of PNPLA3, the genetic underpinning of MASLD-associated bone loss, and provide direction for future studies.

## AUTHOR CONTRIBUTIONS

**GMG** performed the experiments, analyzed the data, and wrote the manuscript. **MS** and **FM** performed experiments. **ACI**, **AHC**, **RL**, and **DCG** assisted in data analysis. **AJS** and **HJD** supervised the studies. All authors have aided in the final preparation of the manuscript and have approved it in its current form.

## SUPPLEMENTAL

### Supplemental Figures

**Supplemental Figure 1:**
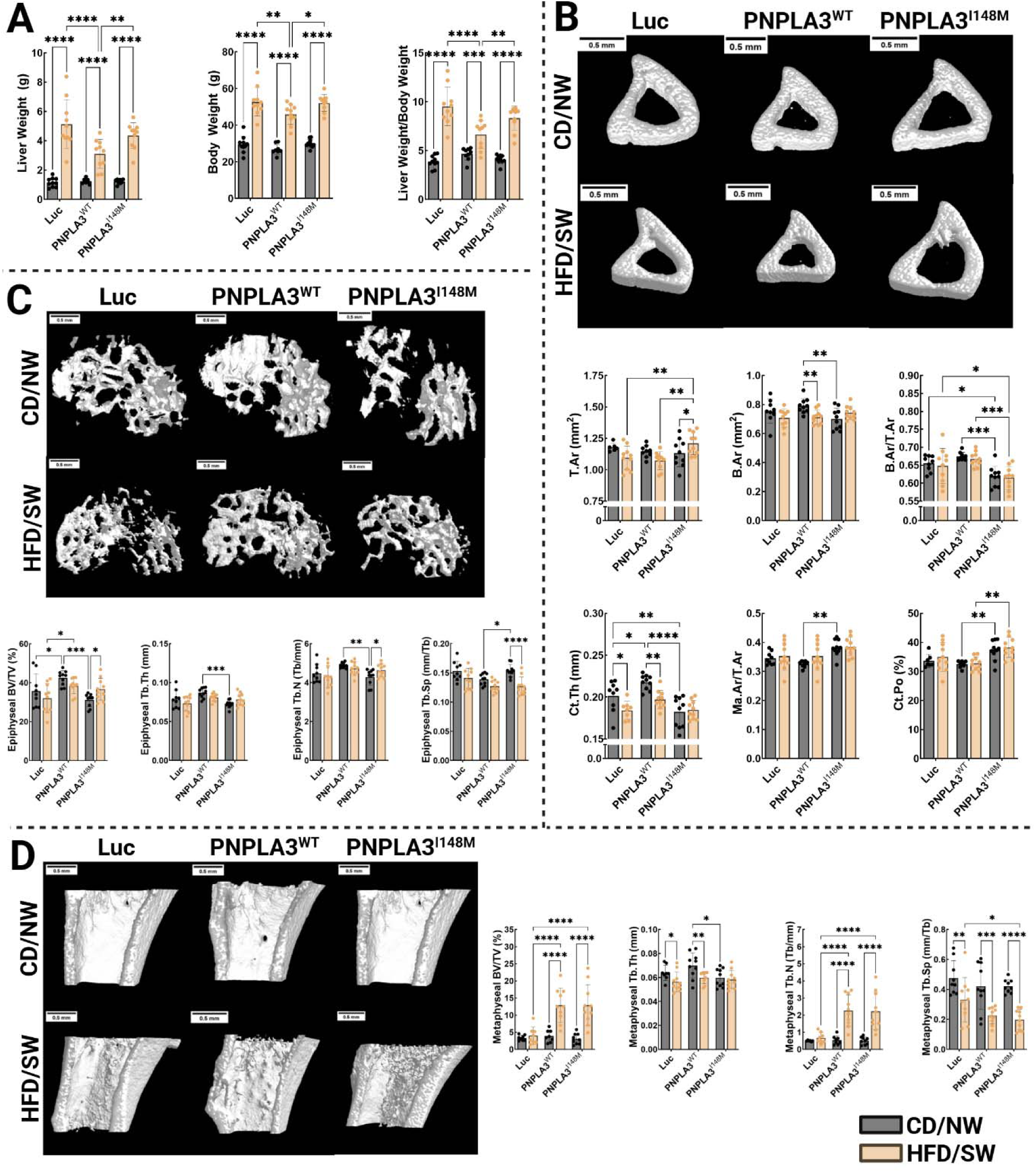
DIAMOND mice fed HFD/SW vs CD/NW become obese, develop hepatomegaly. PNPLA3^WT^ mice lose cortical bone, meanwhile PNPLA3^WT^ and PNPLA3^I148M^ mice gain metaphyseal trabecular bone on HFD/SW vs CD/NW. (**A**) Body and liver weights. 3-dimensional reconstruction and morphological parameters in (**B**) cortical bone, (**C**) epiphyseal trabecular bone, and (**D**) metaphyseal trabecular bone. Data from mice fed CD/NW are also shown in Figure 2. (* p<0.05, ** p<0.01, *** p<0.001, ****p<0.0001)

**Supplemental Figure 2:**
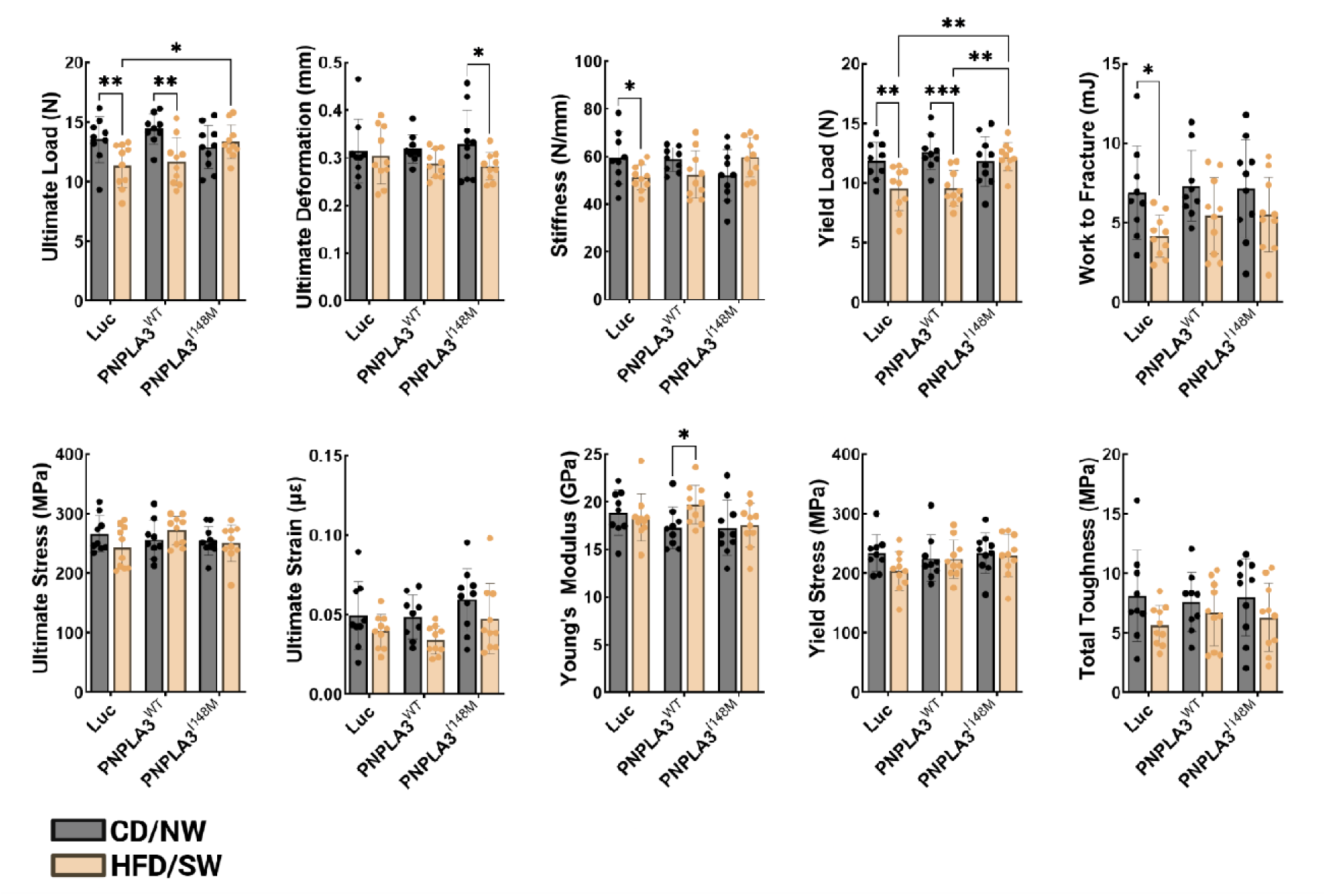
Mechanical testing parameters from *ex vivo* 3-point bending of tibias. Data from mice fed CD/NW are also shown in Figure 2.

**Supplemental Figure 3:**
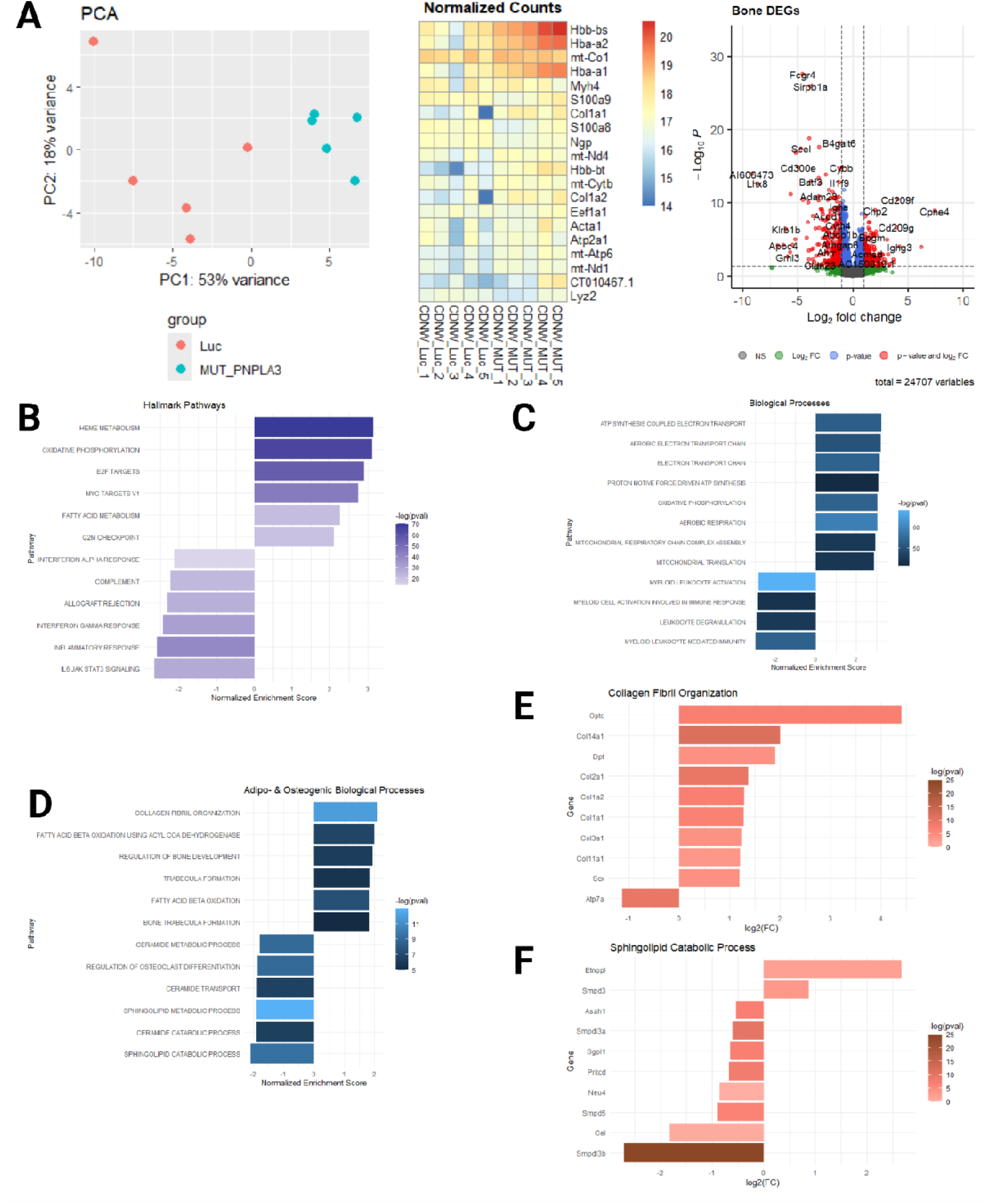
Metabolic, adipogenic, and osteogenic pathways are reprogrammed in bones from PNPLA3^I148M^ mice vs Luc mice. (**A**) Highest expressed genes, separation by PCA, and DEGs in bone from PNPLA3^I148M^ vs Luc mice. (**B**) Hallmark pathway, (**C**) biological process, and (**D**) adipo- and osteogenic ontology pathway enrichment in bone from PNPLA3^I148M^ vs Luc mice. Individual gene contributions to cahnges in (**E**) Collagen fibril organization and (**F**) sphingolipid catabolism biological process ontology terms in bone from PNPLA3^I148M^ mice vs Luc mice.

**Supplemental Figure 4:**
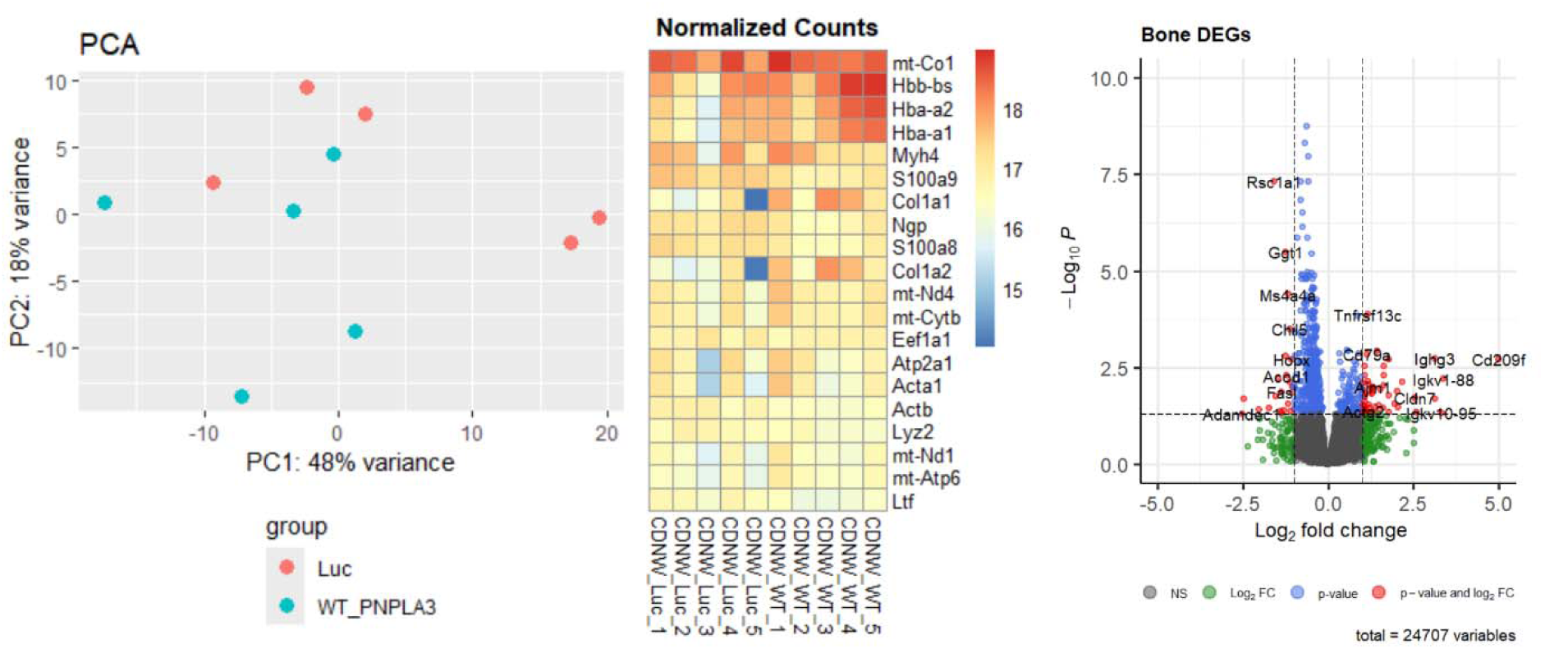
Transcriptomic changes in bone from PNPLA3^WT^ vs Luc mice are sparse. Separation of samples by PCA, highest expressed genes, and differentially expressed genes.

**Supplemental Figure 5:**
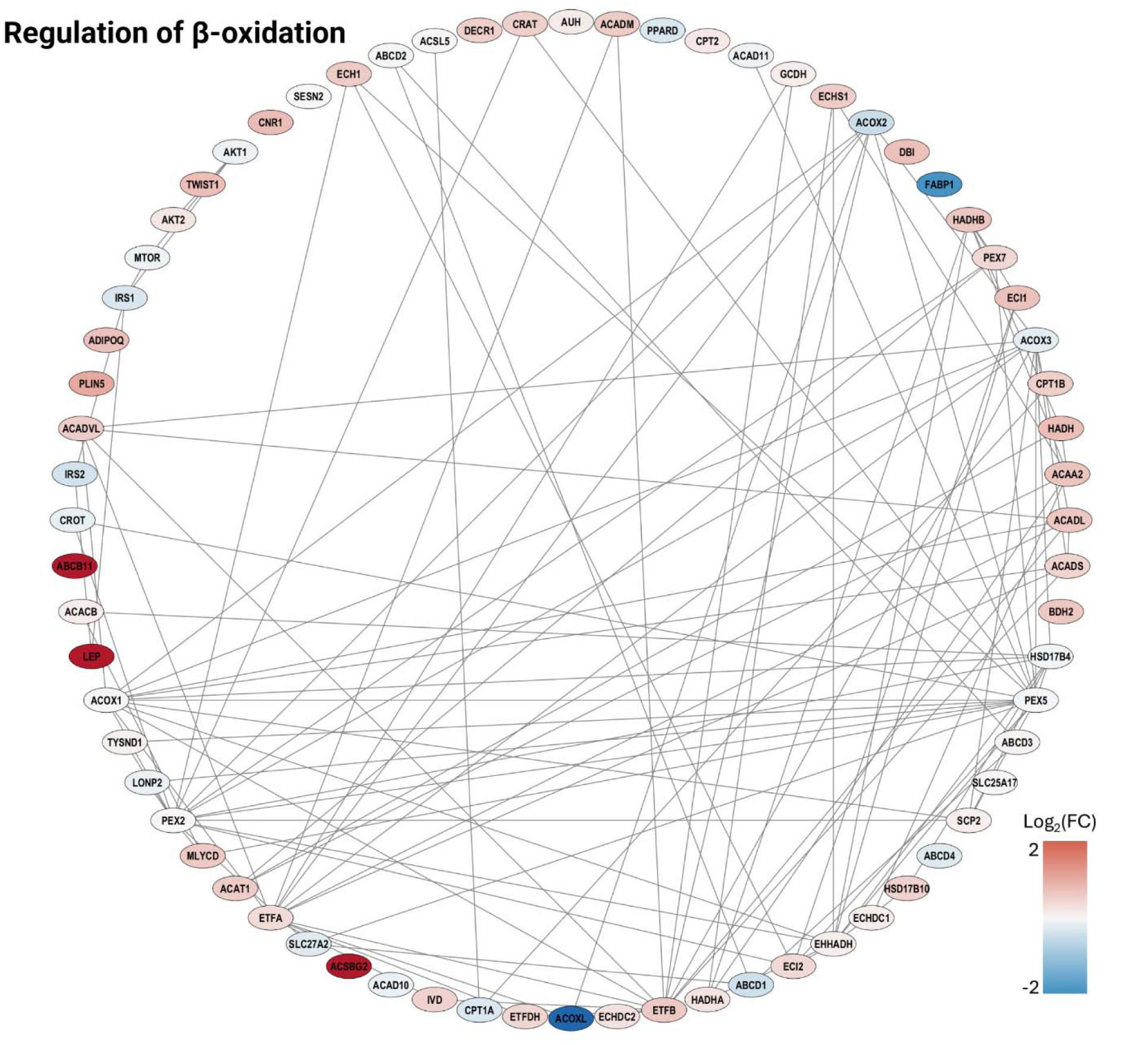
Network regulation of beta oxidation in bone from PNPLA3^I148M^ mice vs PNPLA3^WT^ mice fed CD/NW (inset shown in Figure 4E).

**Supplemental Figure 6:**
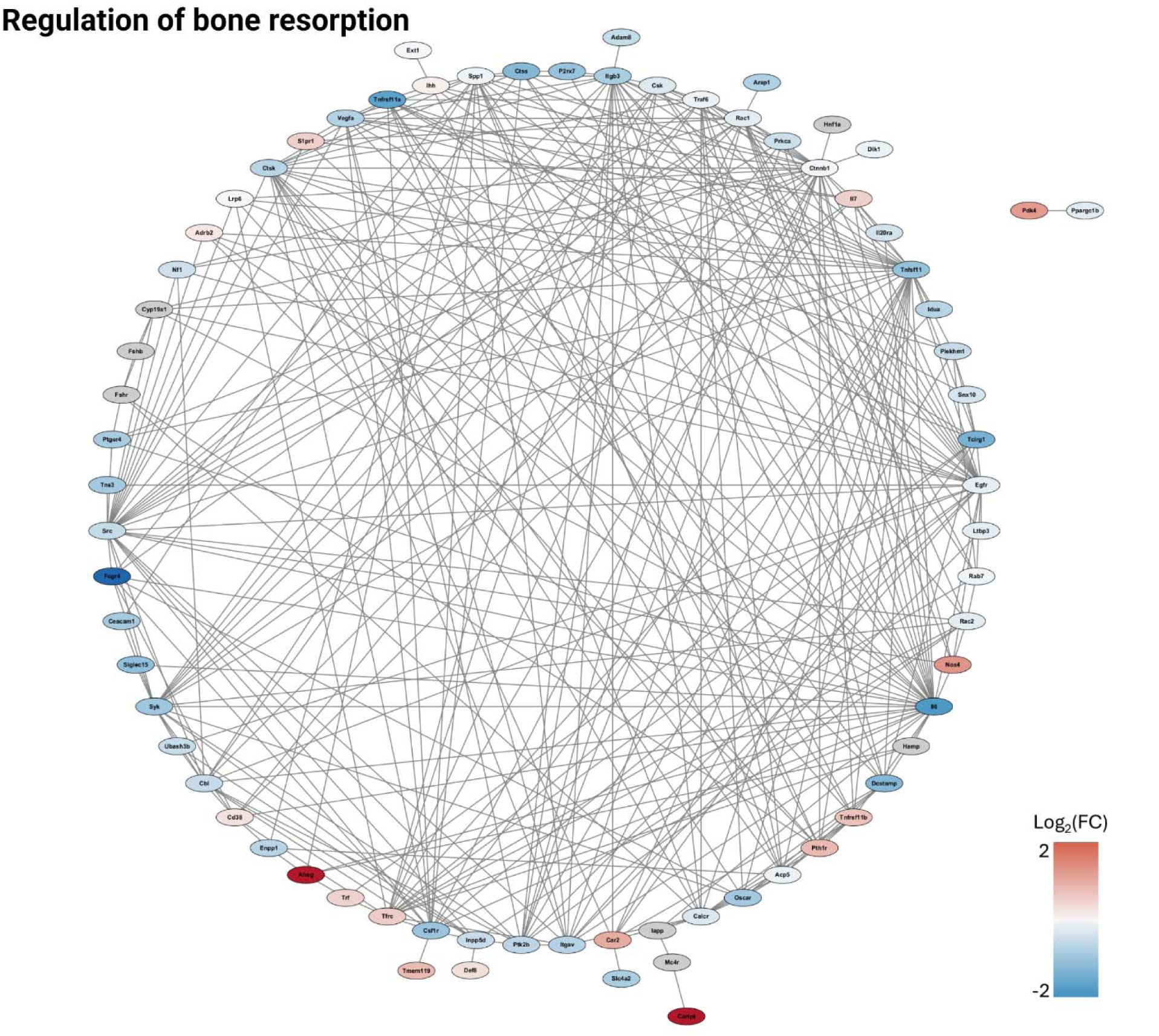
Network regulation of bone resorption in bone from PNPLA3^I148M^ mice vs PNPLA3^WT^ mice fed CD/NW.

**Supplemental Figure 7:**
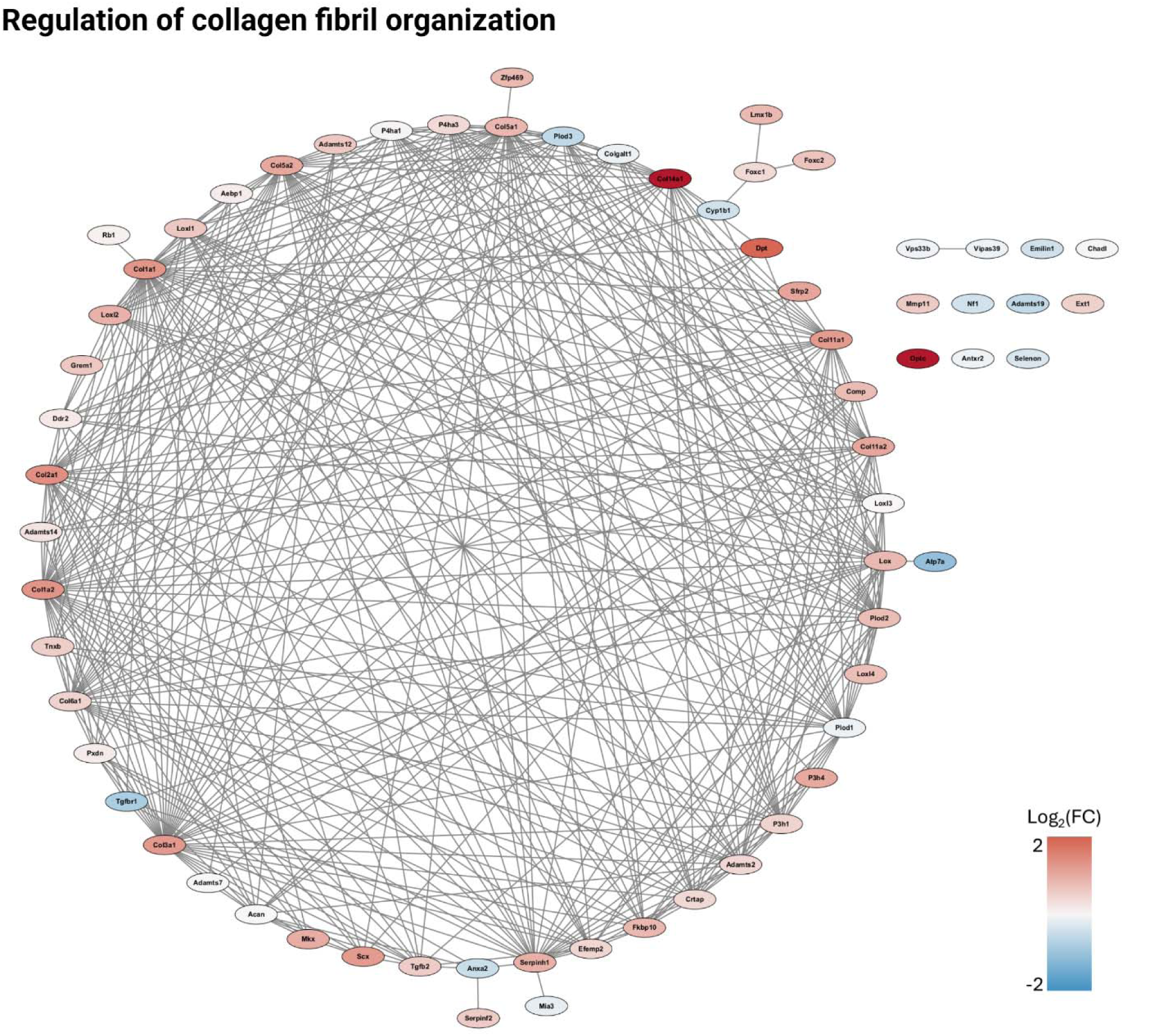
Regulation of collagen fibril organization in bone from PNPLA3^I148M^ mice vs Luc mice fed CD/NW.

**Supplemental Figure 8:**
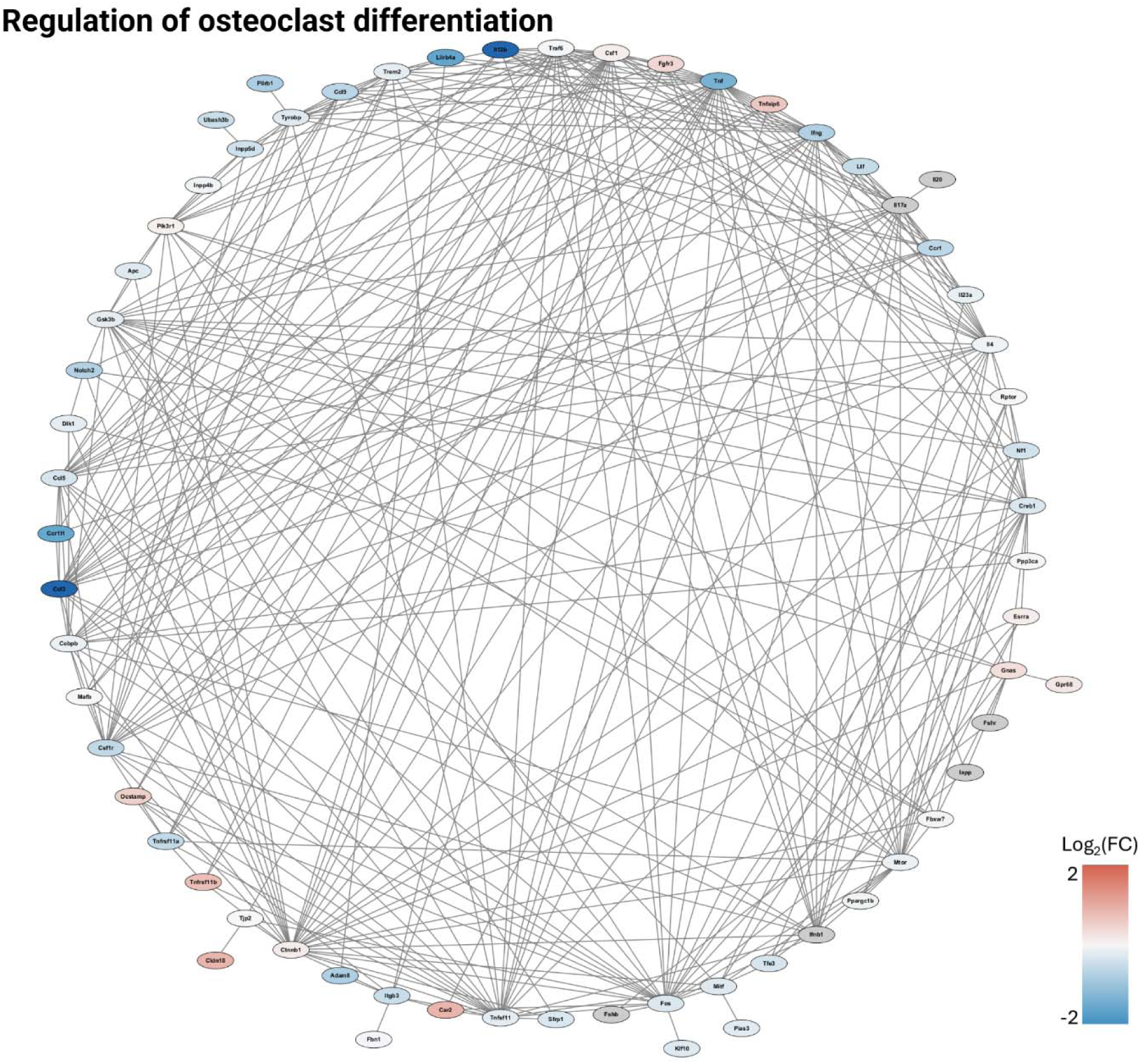
Regulation of collagen fibril organization in bone from PNPLA3^I148M^ mice vs Luc mice fed CD/NW.

## REFERENCES

[1] Z. M. Younossi, M. Kalligeros, and L. Henry, “Epidemiology of metabolic dysfunction-associated steatotic liver disease,” Clin Mol Hepatol, vol. 31, no. Suppl, pp. S32–S50, Aug. 2024, doi: 10.3350/cmh.2024.0431.

[2] A. M. Diehl and C. Day, “Cause, Pathogenesis, and Treatment of Nonalcoholic Steatohepatitis,” New England Journal of Medicine, vol. 377, no. 21, pp. 2063–2072, Nov. 2017, doi: 10.1056/NEJMra1503519.

[3] G. Goldscheitter et al., “MASLD and MASH increase fracture risk in humans and mice by arresting new bone formation,” Res Sq, Nov. 2025, doi: 10.21203/rs.3.rs-7775325/v1.

[4] S. R. Cummings and L. J. Melton, “Epidemiology and outcomes of osteoporotic fractures,” The Lancet, vol. 359, no. 9319, pp. 1761–1767, 2002, doi: 10.1016/S0140-6736(02)08657-9.

[5] S. Romeo et al., “Genetic variation in PNPLA3 confers susceptibility to nonalcoholic fatty liver disease,” Nat Genet, vol. 40, no. 12, pp. 1461–1465, Dec. 2008, doi: 10.1038/NG.257.

[6] Y. Wang et al., “PNPLA3(148M) is a gain-of-function mutation that promotes hepatic steatosis by inhibiting ATGL-mediated triglyceride hydrolysis,” J Hepatol, vol. 82, no. 5, pp. 871–881, 2025, doi: 10.1016/j.jhep.2024.10.048.

[7] Y. Chen et al., “Genome-wide association meta-analysis identifies 17 loci associated with nonalcoholic fatty liver disease,” Nat Genet, vol. 55, no. 10, pp. 1640–1650, 2023, doi: 10.1038/s41588-023-01497-6.

[8] L. I. Plotkin, N. Sanz, and L. R. Brun, “Messages from the Mineral: How Bone Cells Communicate with Other Tissues,” Calcif Tissue Int, vol. 113, no. 1, pp. 39–47, 2023, doi: 10.1007/s00223-023-01091-2.

[9] R. C. Riddle and T. L. Clemens, “Bone Cell Bioenergetics and Skeletal Energy Homeostasis,” Physiol Rev, vol. 97, no. 2, pp. 667–698, Feb. 2017, doi: 10.1152/physrev.00022.2016.

[10] G. M. Goldscheitter et al., “Sexual dimorphism of MASLD-driven bone loss,” Sci Rep, vol. 15, no. 1, p. 23090, 2025, doi: 10.1038/s41598-025-08693-w.

[11] A. Mosca et al., “Relationship between non-alcoholic steatohepatitis, PNPLA3 I148M genotype and bone mineral density in adolescents,” Liver International, vol. 38, no. 12, pp. 2301–2308, Dec. 2018, doi: 10.1111/liv.13955.

[12] R. Xie et al., “Relationship between nonalcoholic fatty liver disease and bone mineral density in adolescents,” Medicine (United States), vol. 101, no. 41, p. E31164, Oct. 2022, doi: 10.1097/MD.0000000000031164.

[13] M. Uhlén et al., “Tissue-based map of the human proteome,” Science (1979), vol. 347, no. 6220, p. 1260419, Jan. 2015, doi: 10.1126/science.1260419.

[14] B. A. Banini et al., “Identification of a Metabolic, Transcriptomic, and Molecular Signature of Patatin-Like Phospholipase Domain Containing 3-Mediated Acceleration of Steatohepatitis,” Hepatology, vol. 73, no. 4, pp. 1290–1306, Apr. 2021, doi: 10.1002/HEP.31609.

[15] C. H. Turner and D. B. Burr, “Basic biomechanical measurements of bone: a tutorial.,” Bone, vol. 14, no. 4, pp. 595–608, 1993, doi: 10.1016/8756-3282(93)90081-k.

[16] P. Bankhead et al., “QuPath: Open source software for digital pathology image analysis,” Sci Rep, vol. 7, no. 1, p. 16878, 2017, doi: 10.1038/s41598-017-17204-5.

[17] A. Cui et al., “Causal association of NAFLD with osteoporosis, fracture and falling risk: a bidirectional Mendelian randomization study.,” Front Endocrinol (Lausanne), vol. 14, p. 1215790, 2023, doi: 10.3389/fendo.2023.1215790.

[18] C. J. Walkey et al., “A comprehensive atlas of AAV tropism in the mouse,” Molecular Therapy, vol. 33, no. 3, pp. 1282–1299, 2025, doi: 10.1016/j.ymthe.2025.01.041.

[19] J. H. Cole and M. C. H. van der Meulen, “Whole Bone Mechanics and Bone Quality,” Clin Orthop Relat Res, vol. 469, no. 8, 2011, [Online]. Available: https://journals.lww.com/clinorthop/fulltext/2011/08000/whole_bone_mechanics_and_bone_quality.6.aspx

[20] S. C. Manolagas, “Birth and Death of Bone Cells: Basic Regulatory Mechanisms and Implications for the Pathogenesis and Treatment of Osteoporosis*,” Endocr Rev, vol. 21, no. 2, pp. 115–137, Apr. 2000, doi: 10.1210/edrv.21.2.0395.

[21] L. M. Tiede-Lewis et al., “Degeneration of the osteocyte network in the C57BL/6 mouse model of aging,” Aging, vol. 9, no. 10, pp. 2190–2208, Oct. 2017, doi: 10.18632/aging.101308.

[22] C. M. Heveran, A. Rauff, K. B. King, R. D. Carpenter, and V. L. Ferguson, “A new open-source tool for measuring 3D osteocyte lacunar geometries from confocal laser scanning microscopy reveals age-related changes to lacunar size and shape in cortical mouse bone,” Bone, vol. 110, pp. 115–127, 2018, doi: 10.1016/j.bone.2018.01.018.

[23] W. Shen et al., “Relationship between MRI-Measured Bone Marrow Adipose Tissue and Hip and Spine Bone Mineral Density in African-American and Caucasian Participants: The CARDIA Study,” J Clin Endocrinol Metab, vol. 97, no. 4, pp. 1337–1346, Apr. 2012, doi: 10.1210/jc.2011-2605.

[24] W. Shen, J. Chen, M. Punyanitya, S. Shapses, S. Heshka, and S. B. Heymsfield, “MRI-measured bone marrow adipose tissue is inversely related to DXA-measured bone mineral in Caucasian women,” Osteoporosis International, vol. 18, no. 5, pp. 641–647, 2007, doi: 10.1007/s00198-006-0285-9.

[25] J. N. Beresford, J. H. Bennett, C. Devlin, P. S. Leboy, and M. E. Owen, “Evidence for an inverse relationship between the differentiation of adipocytic and osteogenic cells in rat marrow stromal cell cultures,” J Cell Sci, vol. 102, no. 2, pp. 341–351, Jun. 1992, doi: 10.1242/jcs.102.2.341.

[26] Y. Wan, L.-W. Chong, and R. M. Evans, “PPAR-γ regulates osteoclastogenesis in mice,” Nat Med, vol. 13, no. 12, pp. 1496–1503, 2007, doi: 10.1038/nm1672.

[27] J. Jules, W. Chen, X. Feng, and Y.-P. Li, “CCAAT/Enhancer-binding Protein α (C/EBPα) Is Important for Osteoclast Differentiation and Activity*,” Journal of Biological Chemistry, vol. 291, no. 31, pp. 16390–16403, 2016, doi: 10.1074/jbc.M115.674598.

[28] J. J. Smink, V. Bégay, T. Schoenmaker, E. Sterneck, T. J. de Vries, and A. Leutz, “Transcription factor C/EBPβ isoform ratio regulates osteoclastogenesis through MafB,” EMBO J, vol. 28, no. 12, pp. 1769–1781, 2009, doi: 10.1038/emboj.2009.127.

[29] A. Bartelt et al., “Quantification of Bone Fatty Acid Metabolism and Its Regulation by Adipocyte Lipoprotein Lipase,” Int J Mol Sci, vol. 18, no. 6, p. 1264, Jun. 2017, doi: 10.3390/ijms18061264.

[30] H. Qiu et al., “Heme metabolism mediates RANKL-induced osteoclastogenesis via mitochondrial oxidative phosphorylation,” Journal of Bone and Mineral Research, vol. 40, no. 5, pp. 639–655, May 2025, doi: 10.1093/jbmr/zjaf040.

[31] I. Palacios-Marin, D. Serra, J. Jimenez-Chillarón, L. Herrero, and M. Todorčević, “Adipose Tissue Dynamics: Cellular and Lipid Turnover in Health and Disease,” Nutrients, vol. 15, no. 18, p. 3968, Sep. 2023, doi: 10.3390/nu15183968.

[32] F. Elefteriou et al., “Serum leptin level is a regulator of bone mass,” Proceedings of the National Academy of Sciences, vol. 101, no. 9, pp. 3258–3263, Mar. 2004, doi: 10.1073/pnas.0308744101.

[33] G. Qian, W. Fan, B. Ahlemeyer, S. Karnati, and E. Baumgart-Vogt, “Peroxisomes in Different Skeletal Cell Types during Intramembranous and Endochondral Ossification and Their Regulation during Osteoblast Differentiation by Distinct Peroxisome Proliferator-Activated Receptors,” PLoS One, vol. 10, no. 12, pp. e0143439-, Dec. 2015, [Online]. Available: 10.1371/journal.pone.0143439

[34] E. F. Eriksen, “Normal and Pathological Remodeling of Human Trabecular Bone: Three Dimensional Reconstruction of the Remodeling Sequence in Normals and in Metabolic Bone Disease*,” Endocr Rev, vol. 7, no. 4, pp. 379–408, Nov. 1986, doi: 10.1210/edrv-7-4-379.

[35] M. von Scheidt et al., “Applications and Limitations of Mouse Models for Understanding Human Atherosclerosis,” Cell Metab, vol. 25, no. 2, pp. 248–261, 2017, doi: 10.1016/j.cmet.2016.11.001.

[36] S. Takahashi et al., “Cyp2c70 is responsible for the species difference in bile acid metabolism between mice and humans[S],” J Lipid Res, vol. 57, no. 12, pp. 2130–2137, 2016, doi: 10.1194/jlr.M071183.

